# Deep RNN with Pseudo Loss Objective for Forecasting Stop-over Decisions of Wild Migratory Birds

**DOI:** 10.1101/2021.04.10.439294

**Authors:** Kehinde Owoeye

## Abstract

Forecasting stop-over decisions and mapping the stop-over sites of wild migratory birds is fast becoming important in light of recent developments affecting global health. Migratory wild birds stop at sites with access to food resources so they can rest before continuing with their journey. Unfortunately, these sites offer opportunities for these birds to spread pathogens and viruses by interacting with the ecosystem. While previous work has focused on predicting stop-over sites using historical information, we emphasize that this is not useful for any planning efforts by health authorities and instead offer a new perspective by proposing an approach that can forecast the duration of stop-over. In this work, first we cast this problem as a classification task and show how pseudo labels and losses in a Bi-directional recurrent neural network can help improve performance given the presence of significantly underrepresented class. We use dataset of Turkey vulture (avian pox vector) movement over several years for the forecasting task where we compare our approach with a variety of baselines and show that it outperforms them. We also use this dataset and the White Fronted Geese (avian flu vector) movement dataset to understand the nature of the habitats used for stop-over using a publicly available model pre-trained on more than half a million land cover images. By knowing the preferred stop-over habitats and the time spent in and between stop-overs using our model, we can help relevant authorities come up with efficient intervention measures.

## I. Introduction

The one health initiative of the World Health Organization aims to control diseases (zoonotic infections) at the interface of human and animal interactions using a multidisciplinary approach, with the recent event affecting global health bringing to the fore again the increasing interdependency in the complex relationships between human and animals. This relationship is enhanced by the encroachment into animal habitats by humans due to developmental activities and climate change. Major human infectious diseases such as Ebola and avian flu have been shown to have animal origin with 20% of this coming from primates alone [1]. The spread of zoonotic infections for instance the highly pathogenic H5N1 avian influenza has huge economic and health implications most importantly for the developing countries of the world. In light of recent developments, it is therefore becoming increasingly imperative that relevant disease surveillance and public health authorities make adequate preparations towards reducing future global health risks using holistic intervention schemes incorporating current and potential relationships between all players in nature’s ecosystem.

A large number of migratory bird species stop-over intermittently to refuel, rest and re-energize [2] for the rest of the long journey as they seek warmer climes for survival during winter and spring migration (also known as flight for survival). The wild variants of these birds such as Geese, Swans, Waterfowls, Eastern wood pewee, Acadian flycatcher, Yellow-green vireo and eagles however are known to be carriers of harmful bacteria, pathogens and viruses from one place to another. These harmful diseases are spread mostly at stop-over sites when these birds interact with resources in the ecosystem such as fresh water via droppings as well as interact with local birds and other non-wild birds using the stop-over sites.

Adequate mapping of sites and forecasting of stop-over decisions thus have potential benefits and could help relevant national and global health authorities come up with several intervention schemes in their response and preparedness measures [3]. While previous works have largely either focused on using historical stop-over sites as a yardstick for predicting future ones [4] using Markov chain or estimate important environmental factors relevant for stopover decisions [5], we point out here that such methods become inadequate given sites that have never been seen before in the training data with the likelihood becoming even higher given climate change activities. Our approach instead integrates this information whilst being flexible enough to potentially predict sites that have never being used before. Also, a direct benefit of our approach is the ability to forecast the duration of the stop-over. Furthermore, we aim to map out the characteristics and features of these sites towards understanding while they are choice for stop-over. Apart from revealing potential stop-over sites in the event of extreme climate change affecting the decision to use historical sites, this can be useful for instance in creating artificial ones in nearby or isolated environments for sustainability purpose and disease containment.

In this work, we consider the task of forecasting the stop-over decisions of migratory birds for the purpose of disease outbreak management, prevention and containment. We cast the task as a binary classification task where we aim to forecast whether an animal is going to move or otherwise. Given the instances where the animals do move are underrepresented in the dataset, we propose a pseudo loss objective function where the original binary cross entropy loss is recast as a multi-class cross entropy loss. We use this approach with the aid of a deep bidirectional recurrent neural network and show that it outperforms several baselines. In addition, we also show using a convolutional neural network model pre-trained on several land cover images, characteristics of these stop-over sites and provide explanations as to what makes these sites desirable and different from non stop-over sites. We evaluate all methods on real world mobility datasets of migratory birds, the former on fine-grained Turkey vulture mobility and the latter on both the Turkey vulture dataset and the White fronted Geese (Anser albifrons) [6], [7] considered to be a wild bird species. The rest of the work is organized as follows, first we review related works followed by a description of the proposed approach and datasets. This is followed by the experiments as well as result and discussion section and we conclude by highlighting some limitations of our work and potential future directions.

## II. Literature Review

The idea of predicting stop-over sites or decisions is relatively new. We build on previous works here while specifying what differentiates our work from previous ones. In addition, we also discuss relevant works in the role of wild birds in spreading infectious diseases and other literatures in the area of avian migration.

Previously, [4] using the dataset of white-fronted Geese collected over multiple years, attempted to forecast the stop-over sites of these birds with the aid of Markov chain using historical information. There are however problems with this approach. First, forecasting stop-over sites that have never been used before becomes difficult and second, estimating duration of stop-over becomes impossible which is useful for planning purposes for instance by health authorities for the purpose of preparedness and response planning. In a related work, [5] used random forest to determine important variables such as roads, ecotone, agricultural land among others relevant for stop-over in migratory whooping crane (*Grus americana*) for conservation purpose and [8] reviewed various methods for estimating stop-over duration in birds. Our work differs in that we are interested in forecasting stop-over decisions using mainly quantitative and not qualitative methods.

With respect to forecasting migration, while [9] recently proposed a recurrent neural network for forecasting avian migration at a finer spatio-temporal granularity, [10] used dataset of bird densities from radar in the Netherlands to predict their migration intensities for applications in improved aviation security. To reduce the deaths of some aquatic animals due to their collision with hydroelectric power plants, [11] proposed a seasonal autoregressive method to forecast their migration intensity in advance using datasets of silver eel migration. While these previous works have tried to predict the migration states/intensity, we are interested in a subset of these which is to forecast the stop-over decisions during spring and autumn migration.

The role of migratory birds in spreading pathogens cannot be overemphasized. Recently, [12] showed the avian influenza virus (AIV) migration rates are higher within flyways than between using a large scale data of gene segments from multiple geographical locations. In a similar vein, [13] used information on phylogenetic associations of virus isolates, migratory bird movements as well as trade in poultry and wild birds to estimate the spread of the highly pathogenic avian influenza in several locations across the world. Also, [14] detected the AIV in wild ducks moving between Europe and Asia using swab samples and fresh droppings. While the approaches described here are mainly in vivo and partially in vitro, ours is mainly in silico. One work that closely resemble ours is the use of serological data with environmental features to model the spread of the West Nile virus in Greece using Geographical Information System technology [15]. Again, our work differs by virtue of the argument already made above in addition to the fact that we are forecasting while they are just predicting. Other models of animal movement behaviour can be found in [16], [17].

## III. Model

Given a time series 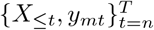 where 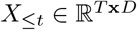 and *y_mt_* is a multi-dimensional vector representing input features (*X*_≤*t*_ = *X_t-d_*,...., *X_t_*, *d* is the duration of the temporal context relevant for the prediction task) and discrete stop-over decision states respectively at each time-step *t* where *m* = 2, the goal is to estimate 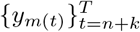 given 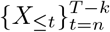 where *k* is the forecast horizon.

### A. Deep Bidirectional RNN

The Bi-directional recurrent neural network [18] learns both backward and forward context for improved sequence modelling compared to conventional RNNs. For increased expressiveness, several layers of this network can be stacked resulting in deep architectures. Given the GRU equations in [19] can be implemented by the function GRU, the deep Bi-GRU for classification with input x, *n* layers and forward 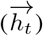 sequence, backward 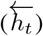 sequence as well as output *y_t_* can then be updated as follows:

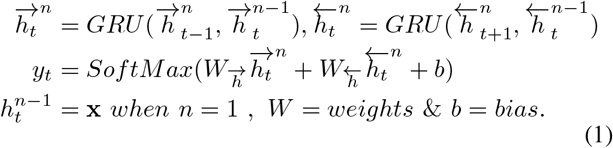

### B. Pseudo Labels, Losses & Training Objectives

A huge number of phenomena in nature are imbalanced or skewed. Using this raw distribution with statistical learning models most times would result in bias outcomes and has generated huge research interest in recent times. One clever approach towards reducing the effect of this imbalance without destroying temporal dependencies inherent in some of these data is to create pseudo labels for the majority class by dividing this class into multiple instances along dimensions that are unique within the class in relation to the input features, whilst being close to the distribution of the underrepresented class as much as possible, but discriminative enough in this regards with respect to the input features. The binary cross entropy loss objective is given by:

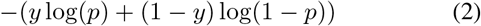

Where *y* represents the class and *p* the observation probability. Instead of minimizing the binary cross entropy objective loss function, given input *X* at training time, we aim to find *θ* that minimizes the overall training objective which is essentially a multi-class cross entropy loss objective within a binary cross entropy loss objective.

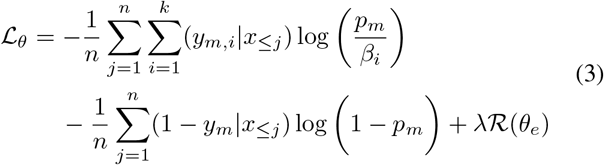

where *y_m,i_* represents the false labels/classes generated from the majority class *y_m_* conditioned on the sequence of input features *x_≤j_* at each data point and the associated probability distribution *p_m_* is divided into smaller segments based on some criterion using *β_i_* (see Algorithm 1 below) which is a function of the length of *y_m,i_* while 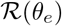 is a regularizer with tuning hyperparameter λ. And the interpretation during training and most especially during testing is such that:

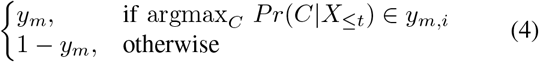

Where *C* represents the new class labels after incorporating pseudo labels such that *y_m,i_* ⊂ *C*.

### C. Connections to Semi-Supervised Learning

The pseudo-label proposed in this work bears some resemblance with the pseudo-label proposed for semi-supervised learning [20] whose loss function is given below:

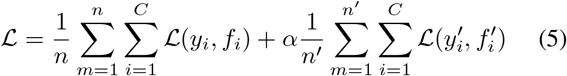

Where *f_i_* is a parameterised function such as a neural network, *C* the number of labels and *y_i_* the labels with the pseudo-label *y*’ defined as:

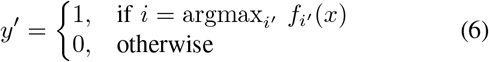

Given that the coefficient balancing term *α*=1, *C* =1 in the first term of the equation, while it is equivalent to the number of pseudo-labels in the second term, and *n*=*n*’, equation (3) is equivalent to (5) when (*y_m,i_*|*x*_≤*j*_) log(*p_m_*) is added to the first term and (1 — *y_m_*|*x*_≤*j*_) log(l — *p_m_*) to the second term of equation (5).

## IV. Datasets & Preprocessing

We evaluate the proposed method on datasets of Turkey vulture described in detail below and also describe the datasets of the White-Fronted Geese.

### A. Turkey vulture & The White-Fronted Geese

The Turkey vulture (*Cathartes aura*) are scavengers with populations spread across North and South America [21]. Turkey vulture movement datasets [22] as part of an ongoing study were collected with the aid of GPS satellite transmitters. We obtain the trajectories of two of the birds (breeding sites in the United States and Canada representative of different populations) with the fewest missing data to expected length of data ratio and with sampling at a resolution of one hour. This bird has been identified as a vector of the avian pox [23]. We use this dataset for the forecasting task due to its fine-grained nature.

The White-Fronted Geese (*Anser albifrons*) on the other hand is a species of the Geese family belonging to a larger group of birds called the waterfowl which include other birds such as Ducks and Swans. Dataset of 65 white-fronted geese [7] migrating between Western Europe and the Russian Arctic were collected over a period of 6 years. This species has been identified as vector of infectious diseases [24] such as the avian influenza virus.

### B. Environmental & Weather Data

The movement trajectories of the Turkey vulture were annotated with weather and environmental data with the aid of the Env-DATA Track Annotation Service [25]. We obtained features such as sunshine duration, snow temperature, population density, atmospheric water, albedo, downward ultraviolet radiation at the surface, plant canopy surface water at surface, incident solar radiation, elevation. dew point temperature, snow evaporation, soil water content, water vapour concentration, snow albedo, temperature parameter in canopy conductance at surface, surface solar radiation downwards, soil temperature, surface solar radiation, maximum temperature, evaporation, ice temperature, land surface temperature night, land surface temperature day, surface thermal radiation and ten metre wind gust.

### C. Satellite Images

Satellite image patches from the Sentinel-2 satellite were collected using customary API. Coordinates corresponding to the first and last seen stop-over points as well as the dates were used as input. We retrieved only the most recent image patches available online relative to the last seen date of the birds at each stopover site each season.

### D. Data Preprocessing

#### 1) Environmental & Weather Data

All missing data in the environmental & weather data were replaced with the last observed points in the sequence. We divided this trajectory (about 28613 in size for the first bird and 11900 for the second) into two, 0.75 of this for training and the rest for testing. The distance (km) covered was computed using the Haversine formula (equation (7)). For the purpose of our task, we consider the movement state as not moving when distance between two contiguous points is less than or equal to 1km and moving if otherwise.

2r arcsin *

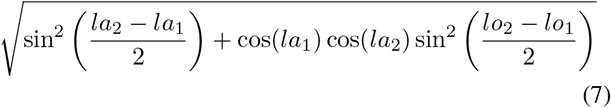

Where r is the radius of earth in kilometers and is equal to 6371 and *la*_2_,*la*_1_ represents contiguous latitude points while *lo*_2_,*lo*_1_ represents contiguous longitude points.

#### 2) Images

The downloaded image patches were atmospherically corrected. The sentinel-2 images have 13 spectral bands spread across three resolutions of 10m, 20m and 60m. The images were down sampled (10m band: 120×120 pixels, 20m band: 60×60 pixels and 60m band: 20×20 pixels) as in [26] and converted from JPEG to TIF. We removed one of each duplicated image corresponding to instances where the bird left and returned to the same stop-over site.

## V. Experiments & Procedures

We evaluate all models using accuracy and F1 scores of all the test data and the migration window alone. All deep learning models were trained thrice and evaluated on the test set for the same amount of time. We report the mean and standard deviations of these results. To recast the problem as a multiclassification task, for simplicity, we divide the majority class along the dimension of migration or movement state. Here, there are three movement states corresponding to breeding, wintering and migration where the fall and spring migration states have been merged together (see Algorithm 1 below where *y_mt_* and *M^it^* represents the mobility and migration states respectively.

### A. Model Architecture & Parameters

We use a four layer bidirectional recurrent neural network scanning in both directions implemented using the gated recurrent unit [19] where the input representing the data described in section IV (C) and the movement coordinates are fed to the network using a many to one architecture where the output represents the pseudo labels. The hyper-parameters used include the Adam optimizer [27], cell units = 60, time step = 60, batch size = 100 learning rate = 1e-2 and a SoftMax at the output layer.

### B. Experiments

We design several experiments described in detail below to answer several questions that can help shed light with respect to the objectives of this work.

#### 1) Turkey vulture, Q1

How does the proposed approach perform compared to some baselines? To investigate this, we use the proposed approach and compare the performance with several baselines described below. Forecast horizon of up to 24hrs are considered as the birds don’t usually spend up to a day at each stop-over location.

#### 2) Turkey vulture, Q2

How does the proposed approach perform when used without the pseudo labels? We use this as an ablation and compare the proposed model with one trained using a binary cross entropy loss.

#### 3) Turkey vulture, Q3

How do the baseline models perform even when used with pseudo labels? We investigate if the proposed loss objective can improve the performance of the baseline models.

#### 4) Turkey vulture, Q4

How do classical techniques for handling imbalanced classification technique perform on sequential data compared to our approach? We compare techniques such as under-sampling with the proposed approach for instance we set class weight to balanced for random forest and logistic regression and use weighted class entropy losses for the deep learning models.

#### 5) Turkey vulture, Q5

Can we estimate the duration of stop-over and how does this compare with the ground truth? To answer this question, we compare the results of the model relative to the ground truth at each stopover period greater than or equal to 2 hours relative to the ground truth data.

#### 6) Turkey vulture, Q6

While the nature of the breeding and wintering habitats of this bird is known, we ask, what are the peculiar features of the Turkey vulture stop-over sites? To answer this question, we extract instances where there has been significant time spent during stop-overs in the dataset. See Q7 below for detailed description of the process involved. We use the bird with the biggest movement trajectory here.

#### 7) White Fronted Geese, Q7

What are the features peculiar to the stop-over sites of these birds? We use the pre-trained ResNet-50 architecture with a fully connected layer at the output consisting of 19 neurons (model of [26]). The labels/classes are provided from the CORINE Land Cover database of the year 2018 (CLC 2018) and merged into 19 groups for easy classification. Since this is a multi-label classification task, we select a probability threshold ≥ 0.1 to observe what land cover/habitats dominates in the image patches. We validate the output of this model by crosschecking it with some of the coordinates that are well known locations/places online (including Wikipedia). We use the bird with the biggest movement trajectory per year over a period of five years.

### C. Baselines

We compare our approach to several baselines including deep and classical learning methods as well as carried out some ablation experiments.

#### 1) Markov Chain (MC)

Markov chain was used in [4] to forecast stop-over sites but this time around to forecast stopover decisions. Here, we use only the labels (the decision to move or not) as input to the model.

**Algorithm 1.**
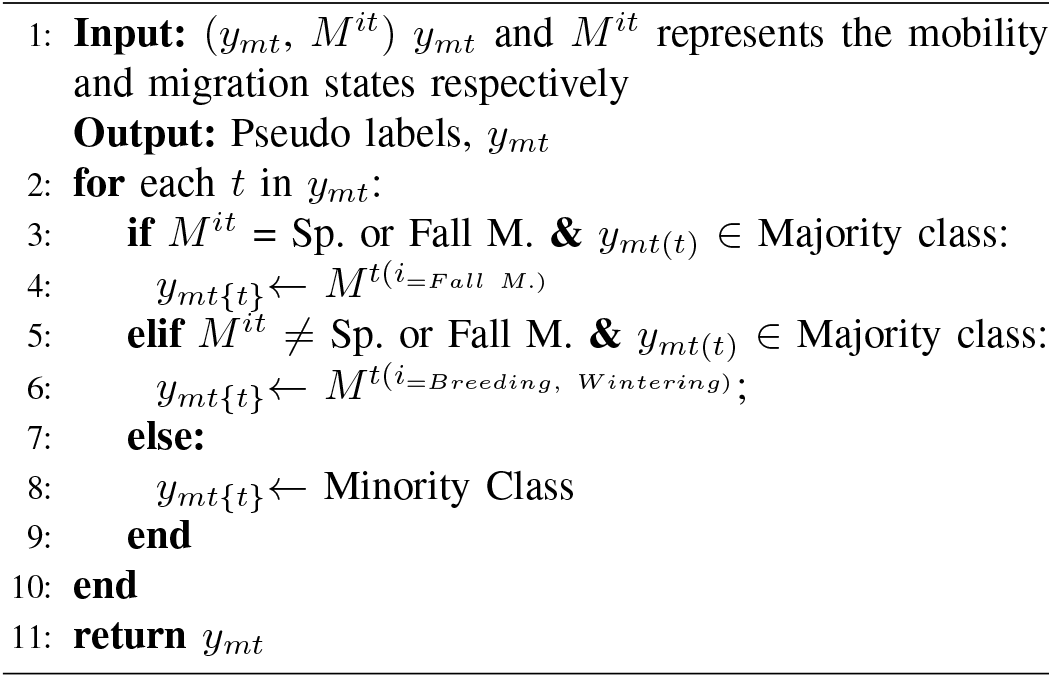
Generating Pseudo labels

#### 2) Deep Bidirectional Recurrent Neural Network with Auxiliary Task (DBi-RNN 2-L, A. Task)

A deep bidirectional recurrent network for forecasting migration states augmented with a mean square error loss by forecasting the longitude [9].

#### 3) Deep Neural Network (DNN)

With two layers and 256 neurons each, dropout = 0.2, SoftMax layer and Adam optimizer [27]. We train all neural networks given similar procedures described for our approach above.

#### 4) Recurrent Neural Network (RNN)

A GRU network and also a bidirectional GRU network with 1 layer, 50 cells, dropout = 0.2, SoftMax layer and Adam optimizer [27].

#### 5) Logistic Regression (LR), Support Vector Machine (SVM) & Random Forest (RF)

We use a multinomial variant with a lbfgs solver for logistic regression and one versus one strategy for SVM. We also use random forest as it was used in [5] to determine what environmental factors are important for stopover decisions.

## VI. Results & Discussion

We discuss the result of the experiments as well as the answers to the questions asked above here in the order listed.

### 1) A1

Results in Table II show the proposed approach perform better than a variety of baselines. Not only are the accuracies better but the F1 scores for the minority class across the experiments are significantly better.

**TABLE I.** Distribution of the two classes of interest for the two birds showing the decision to move class is in the minority.

| Data | Move | N. Moving |
| --- | --- | --- |
| Training | 0.22, 0.21 | 0.78, 0.79 |
| Test | 0.23, 0.15 | 0.77, 0.85 |

**TABLE II.**
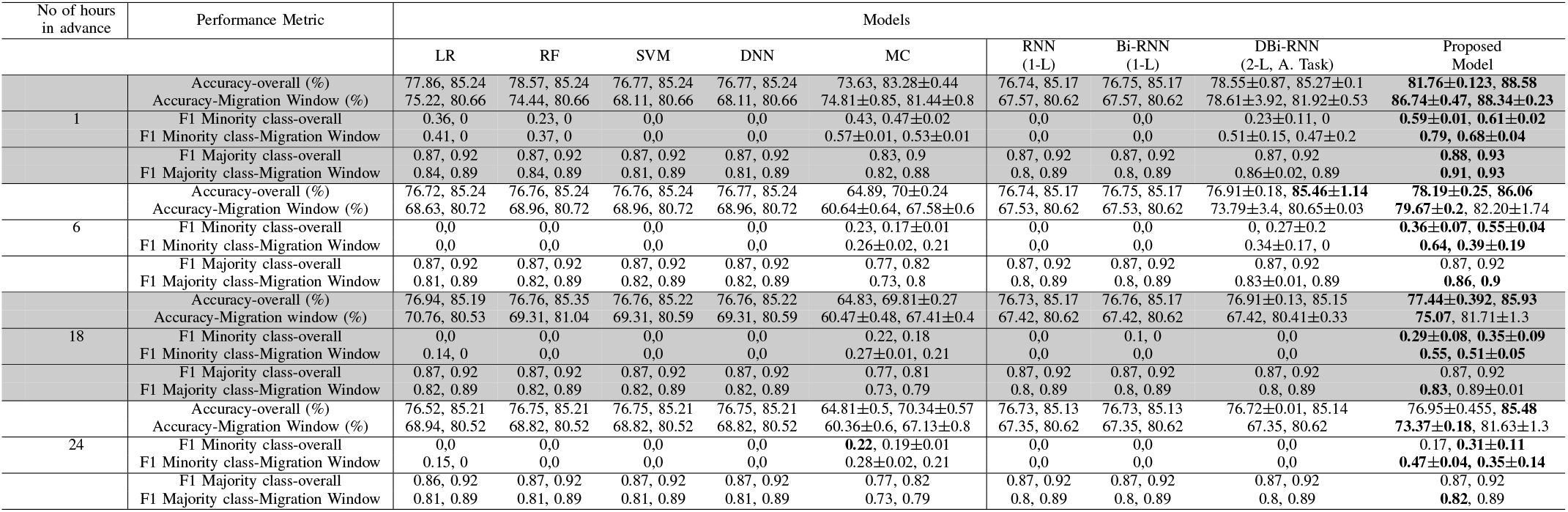
Performance comparison. Our approach can be seen to outperform all baselines across the four experiments. Confidence intervals omitted when less than 0.01, 1-L means one layer while the two figures per entry represent the results for the two birds considered in this work in the order of the biggest to the smallest trajectory size. Bold numbers correspond to the best performing model.

### 2) A2

Results in Table III show that without the pseudo loss, the performance of the proposed model degrades slightly suggesting the effectiveness of the proposed approach.

**TABLE III.**
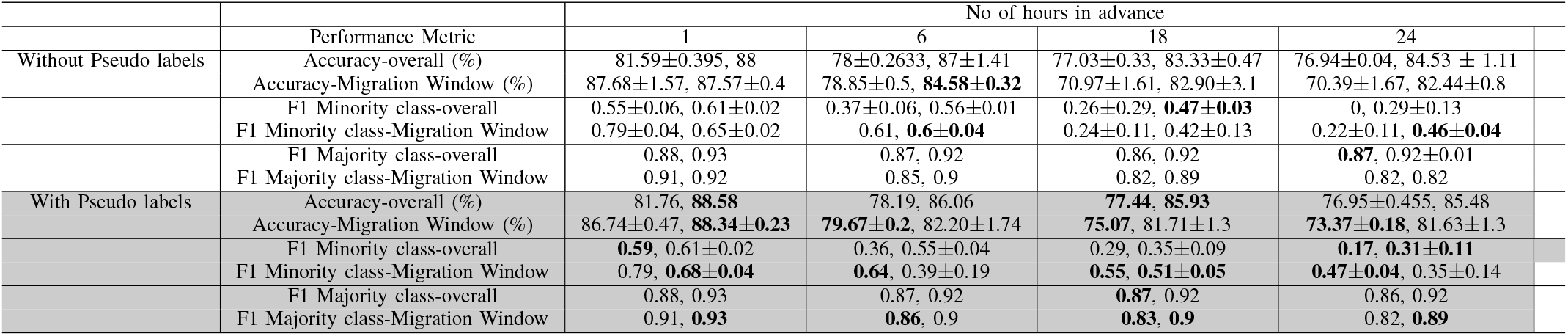
Performance of model with and without pseudo loss objectives. The performance with the pseudo loss objectives can be seen to be better most of the time compared to without it. Confidence intervals omitted when less than 0.01 while the two figures per entry represent the results for the two birds considered in this work in the order of the biggest to the smallest trajectory size. Bold numbers correspond to the best performing model.

### 3) A3

Results in Table IV show the performance of the baselines degrades slightly on the average when used with the pseudo labels but the F1 scores for the minority class improved most especially for the deep learning sequence models. This demonstrates to some extent the generalisation of the proposed approach to other deep sequence architectures.

**TABLE IV.**
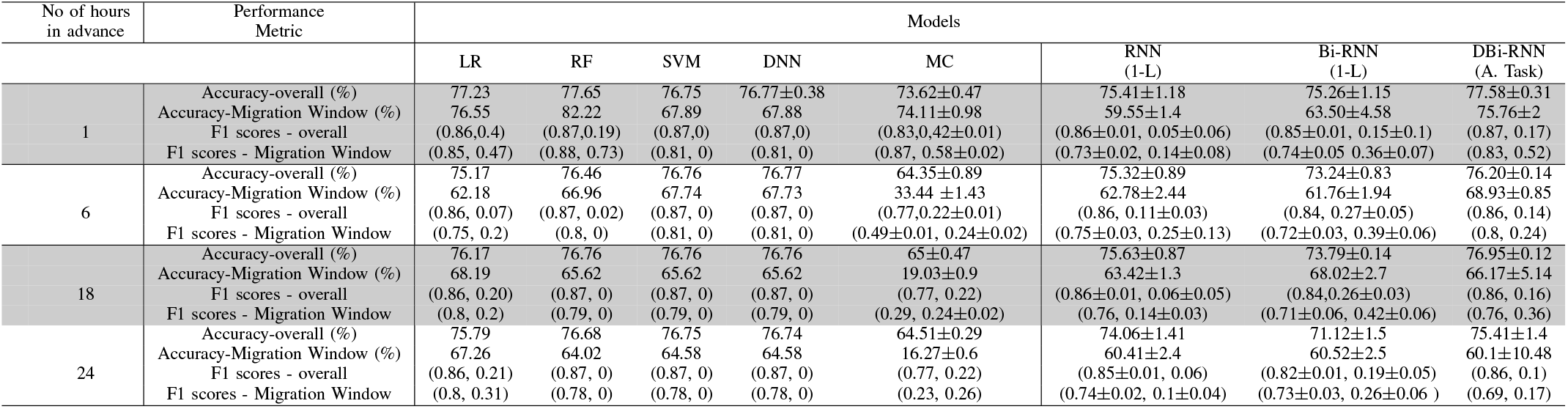
Performance comparison of Baselines with Pseudo labels. Overall, the performance of the baseline models can be seen on the average to have degraded slightly when used with pseudo labels. Only results for the bird with the biggest trajectory have been reported here for brevity and the F1 scores have been reported with respect to the majority class first.

| No of hours<br>in advance | Performance<br>Metric | Models |  |  |  |  |  |  |  |
| --- | --- | --- | --- | --- | --- | --- | --- | --- | --- |
|  |  | LR | RF | SVM | DNN | MC | RNN<br>(1-L) | Bi-RNN<br>(1-L) | DBi-RNN<br>(A. Task) |
| 1 | Accuracy-overall (%) | 77.23 | 77.65 | 76.75 | 76.77±0.38 | 73.62±0.47 | 75.41±1.18 | 75.26±1.15 | 77.58±0.31 |
|  | Accuracy-Migration Window (%) | 76.55 | 82.22 | 67.89 | 67.88 | 74.11±0.98 | 59.55±1.4 | 63.50±4.58 | 75.76±2 |
|  | F1 scores - overall | (0.86,0.4) | (0.87,0.19) | (0.87,0) | (0.87,0) | (0.83,0.42±0.01) | (0.86±0.01, 0.05±0.06) | (0.85±0.01, 0.15±0.1) | (0.87, 0.17) |
|  | F1 scores - Migration Window | (0.85, 0.47) | (0.88, 0.73) | (0.81, 0) | (0.81, 0) | (0.87, 0.58±0.02) | (0.73±0.02, 0.14±0.08) | (0.74±0.05 0.36±0.07) | (0.83, 0.52) |
| 6 | Accuracy-overall (%) | 75.17 | 76.46 | 76.76 | 76.77 | 64.35±0.89 | 75.32±0.89 | 73.24±0.83 | 76.20±0.14 |
|  | Accuracy-Migration Window (%) | 62.18 | 66.96 | 67.74 | 67.73 | 33.44 ±1.43 | 62.78±2.44 | 61.76±1.94 | 68.93±0.85 |
|  | F1 scores - overall | (0.86, 0.07) | (0.87, 0.02) | (0.87, 0) | (0.87, 0) | (0.77,0.22±0.01) | (0.86, 0.11±0.03) | (0.84, 0.27±0.05) | (0.86, 0.14) |
|  | F1 scores - Migration Window | (0.75, 0.2) | (0.8, 0) | (0.81, 0) | (0.81, 0) | (0.49±0.01, 0.24±0.02) | (0.75±0.03, 0.25±0.13) | (0.72±0.03, 0.39±0.06) | (0.8, 0.24) |
| 18 | Accuracy-overall (%) | 76.17 | 76.76 | 76.76 | 76.76 | 65±0.47 | 75.63±0.87 | 73.79±0.14 | 76.95±0.12 |
|  | Accuracy-Migration Window (%) | 68.19 | 65.62 | 65.62 | 65.62 | 19.03±0.9 | 63.42±1.3 | 68.02±2.7 | 66.17±5.14 |
|  | F1 scores - overall | (0.86, 0.20) | (0.87, 0) | (0.87, 0) | (0.87, 0) | (0.77, 0.22) | (0.86±0.01, 0.06±0.05) | (0.84,0.26±0.03) | (0.86, 0.16) |
|  | F1 scores - Migration Window | (0.8, 0.2) | (0.79, 0) | (0.79, 0) | (0.79, 0) | (0.29, 0.24±0.02) | (0.76, 0.14±0.03) | (0.71±0.06, 0.42±0.06) | (0.76, 0.36) |
| 24 | Accuracy-overall (%) | 75.79 | 76.68 | 76.75 | 76.74 | 64.51±0.29 | 74.06±1.41 | 71.12±1.5 | 75.41±1.4 |
|  | Accuracy-Migration Window (%) | 67.26 | 64.02 | 64.58 | 64.58 | 16.27±0.6 | 60.41±2.4 | 60.52±2.5 | 60.1±10.48 |
|  | F1 scores - overall | (0.86, 0.21) | (0.87, 0) | (0.87, 0) | (0.87, 0) | (0.77, 0.22) | (0.85±0.01, 0.06) | (0.82±0.01, 0.19±0.05) | (0.86, 0.1) |
|  | F1 scores - Migration Window | (0.8, 0.31) | (0.78, 0) | (0.78, 0) | (0.78, 0) | (0.23, 0.26) | (0.74±0.02, 0.1±0.04) | (0.73±0.03, 0.26±0.06 ) | (0.69, 0.17) |

### 4) A4

Results in Table V show that, these techniques on average hurts the performance with respect to accuracy but enhanced the performance of the F1 score for the underrepresented class at the expense of the majority class while the proposed approach is able to improve the classification of the minority class without hurting the majority. In most cases also, these techniques destroy the temporal dependency in the dataset giving rise to results not monotonic.

**TABLE V.** Performance of baseline models with other data imbalance techniques. Some of the results are not monotonic for reasons we suspect to be the destruction of the temporal dependency in the data by the use of special resampling and re-weighting techniques thus rendering these methods not so useful for tasks such as forecasting. Only results for the bird with the biggest trajectory have been reported here for brevity and the F1 scores have been reported with respect to the majority class first.

| No of hours<br>in advance | Performance<br>Metric | Models |  |  |  |  |  |
| --- | --- | --- | --- | --- | --- | --- | --- |
|  |  | LR | RF | DNN | RNN<br>(1-L) | Bi-RNN<br>(1-L) | Bi-RNN<br>(4-L) |
| 1 | Accuracy-overall (%) | 75.98 | 75.37 | 76.77 | 24.35 | 32.82 | 79± 0.01 |
|  | Accuracy-Migration Window (%) | 83.11 | 85.11 | 73.40±7.49 | 32.09±0.4 | 55.981.1 | 84.22±2.3 |
|  | F1 scores - overall | (0.83,0.57) | (0.82,0.59) | (0.87,0) | (0, 0.372) | (0.22±0.1,0.4±0.01) | (0.85±0.01, 0.65±0.01) |
|  | F1 scores - Migration Window | (0.88, 0.72) | (0.89, 0.77) | (0.83±0.03, 0.26) | (0, 0.48±0.01) | (0.35±0.22, 0.54±0.03) | (0.87±0.02, 0.79±0.02) |
| 6 | Accuracy-overall (%) | 56.13 | 56.08 | 76.74 | 24.87 | 50.81±0.1 | 70±0.02 |
|  | Accuracy-Migration Window (%) | 55.28 | 53.61 | 68.96 | 34.5±1.9 | 55.78±2.46 | 74.62±3 |
|  | F1 scores - overall | (0.64,0.44) | (0.63, 0.46) | (0.87,0) | (0,0.376) | (0.78± 0.02,0.45± 0.03) | (0.54± 0.16,0.56±0.01) |
|  | F1 scores - Migration Window | (0.58, 0.52) | (0.51, 0.56) | (0.82, 0) | (0, 0.49) | (0.55±0.05, 0.56±0.01) | (0.78±0.03, 0.7±0.02) |
| 18 | Accuracy-overall (%) | 67.57 | 63.70 | 76.75 | 23.59 | 54.15±0.1 | 63.77± 0.014 |
|  | Accuracy-Migration Window (%) | 67.41 | 65.96 | 68.27±1.47 | 32.92±0.4 | 48.77±11.75 | 62.88±7.51 |
|  | F1 scores - overall | (0.77,0.44) | (0.72,0.46) | (0.87,0) | (0,0.376) | (0.6±0.11,0.44±0.02) | (0.71±0.02,0.5) |
|  | F1 scores - Migration Window | (0.75, 0.52) | (0.73, 0.53) | (0.81±0.01, 0) | (0, 0.49) | (0.31±0.28, 0.52±0.03) | (0.64±0.1, 0.61±0.04) |
| 24 | Accuracy-overall (%) | 67.22 | 66.64 | 76.75 | 23.34±0.1 | 23.51±0.27 | 57.6±4.2 |
|  | Accuracy-Migration Window (%) | 70.95 | 72.18 | 68.82 | 32.61±0.4 | 33.41±1.07 | 64.35±1.42 |
|  | F1 scores - overall | (0.77, 0.42) | (0.76, 0.44) | (0.87, 0) | (0, 0.38) | (0, 0.37) | (0.65, 0.46) |
|  | F1 scores - Migration Window | (0.78, 0.55) | (0.79, 0.57) | (0.81, 0) | (0, 0.49) | (0, 0.48±0.02) | (0.66±0.02, 0.62) |

### 5) A5

Results in Fig 4 show the duration of stopover can be estimated with accuracy which appears to decrease as the forecast horizon increases and on the average higher for fall migration stop-overs. In addition, the accuracy is higher for the fall migration on the average because fall migration is generally longer than spring migration meaning more training data and more confidence in the estimation for fall relative to spring stop-overs.

**Fig. 1.**
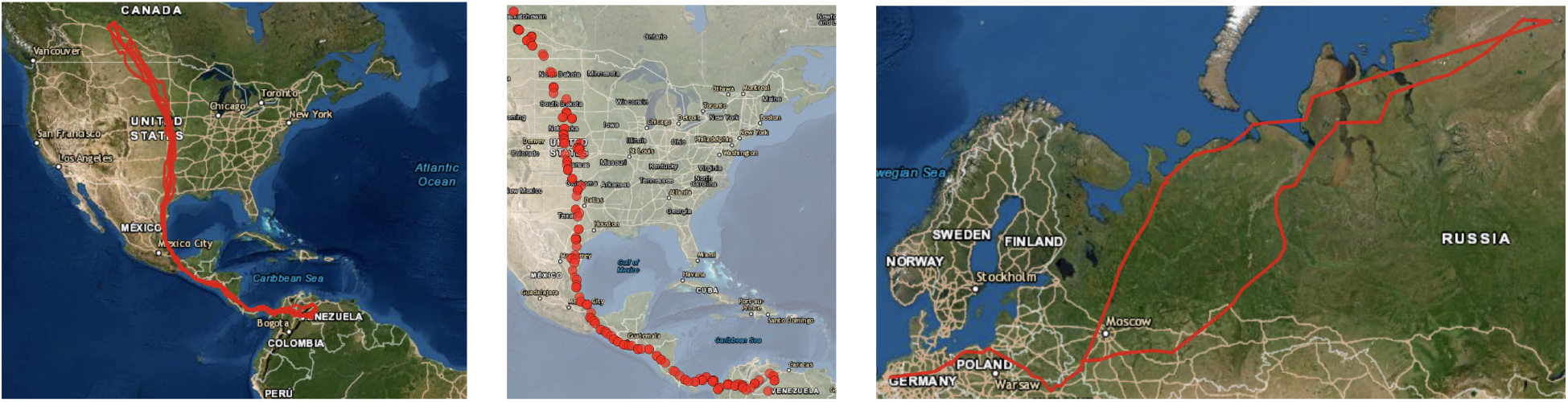
Left: Trajectory of the Turkey vulture as it moves from its breeding site in north America to its wintering site in South America. Middle: Waypoints (stop-over) sites of the Turkey vulture. Right: Trajectory of the White Fronted Geese as they move between their wintering site in the Netherlands to the breeding site in the Russian Arctic.

**Fig. 2.**
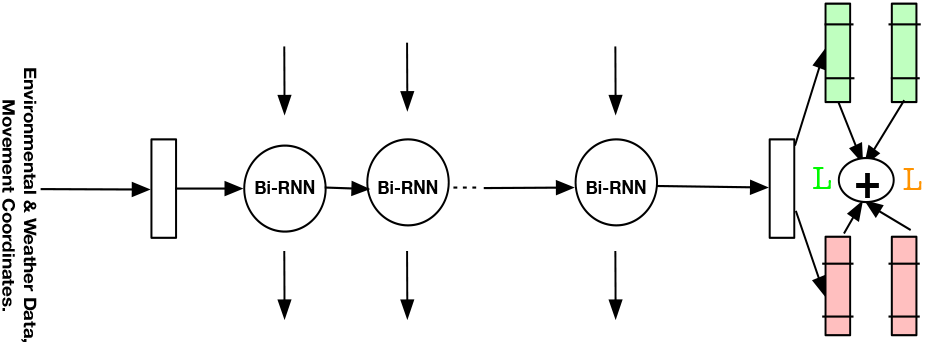
Schematic of the proposed architecture. The white rectangles represent fully connected/input layers while the green & red ones represent respectively the model predictions & targets for the majority class and the pseudo labels. The losses 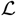 are then summed up and backpropagated to learn the weights of the network.

**Fig. 3.**
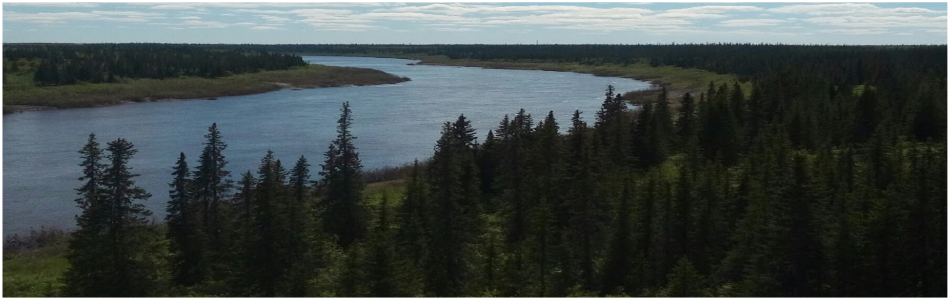
Typical stopover site. Mixture of forest with cone shaped trees and water as seen via google map. The White Fronted Geese and Turkey vulture stop at sites predominantly with access to water.

**Fig. 4.**
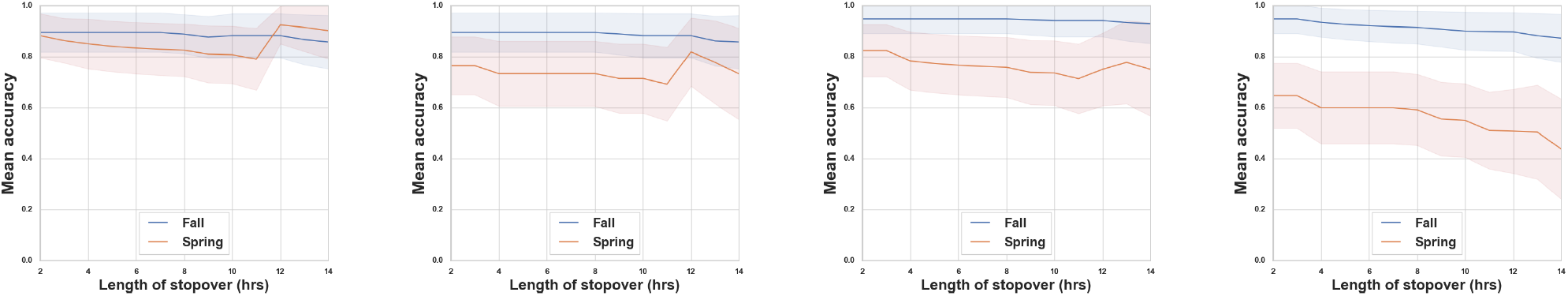
Duration of Stopover. Left to Right: Forecast horizon of 1, 6, 18 and 24 hours respectively. The *y* axis represents the mean proportion of the stopover duration on the x-axis that has been estimated correctly up until the first decision to move is predicted relative to the ground truth, with zero being the lowest and 1 the highest. The bird with the biggest trajectory has been used here as it has both spring and fall migration information in the test data.

### 6) A6

Results show this bird stop-over at places with access to water (for brevity, since the result here is similar to that of the White Fronted Geese, see A7 below for more explanation and figures). While it is surprising this bird never stopped over in stereotype sites with access to carrions, Turkey vultures are known to feed on insects and washed-up fishes [28] and this could be a strategy to have access to both food and water simultaneously with little effort or the priority could just be on water only as some birds eat enough to sustain them before embarking on migration.

### 7) A7

One particular feature common to these sites is the presence of water (Fig 5). This makes sense given the affinity of this avian species for water. The risks with respect to the spread of avian flu however are via droppings and interactions at water bodies in urban areas or areas with high population density along migration path most especially when such water bodies are used for domestic and recreational activities.

**Fig. 5.**
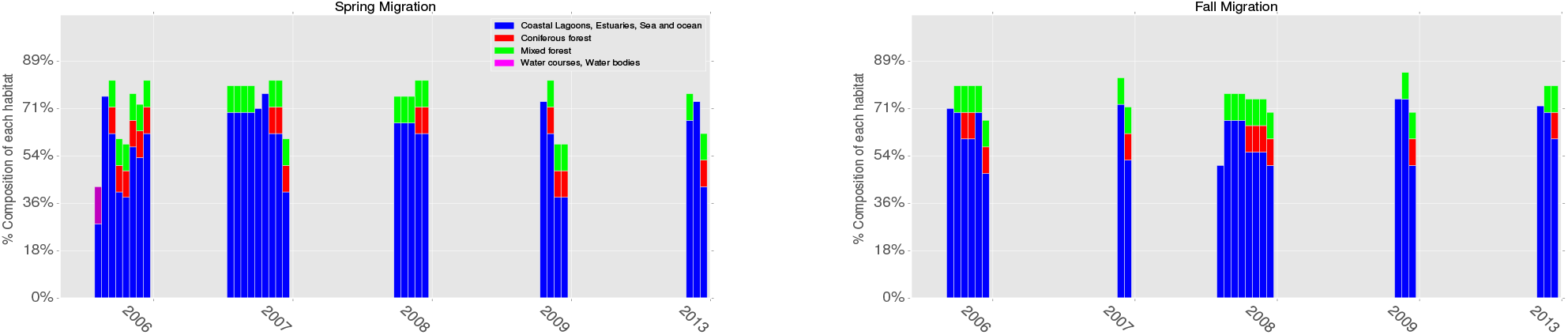
Mapping of habitat. Each bar chart represents a stop-over period where all the constituent groups in each bar sum up to 100% and only habitat with probability ≥ 0.1(10%) are reported. As seen, the birds stop-over at sites with access to water. Note that Turkey vultures stop-over at similar habitats but we omit the figure for brevity.

## VII. Conclusion

We have proposed a deep bidirectional recurrent neural network with pseudo loss objective for forecasting mobility decisions of migratory birds. We also mapped out the preferred stop-over habitats using a pre-trained CNN model. our work however is without its limitations. First, we have ignored the prediction of movement velocities here for simplicity due to the quality and quantity of data used. We aim to incorporate this in our model in the future as more fine-grained data collected over long period becomes available. The model used for mapping the habitats is by no means perfect but good enough to give a high level and general description of the land cover in the image patches without much ambiguity. Additional and relevant environmental and weather features will enable us to consider longer forecast horizons. To achieve this, we aim to look in depth on what factors influence the duration of the stop-overs in future work. Beyond global health concerns, our work also has implications for worldwide conservation efforts and we aim to consider variations of the problem addressed here in this regard in the future.

